# A ‘Variant Viewer’ for investigating population-level variation in *Arabidopsis thaliana*

**DOI:** 10.1101/2021.04.21.440793

**Authors:** Matthew Cumming, Eddi Esteban, Vincent Lau, Asher Pasha, Nicholas J. Provart

## Abstract

High throughput sequencing has opened the doors for investigators to probe genetic variation present in large populations of organisms. In plants, the 1001 Genomes Project (1001genomes.org) is one such effort that sought to characterize the extant worldwide variation in *Arabidopsis thaliana* for future analyses to compare and draw upon. We developed a web application that accesses the 1001 Genomes database called The Variant Viewer, for investigators to view variants in any *A. thaliana* gene and within gene families. These variants may be visualized in the context of alignments of queried genes, across splice isoforms of these genes and in relation to conserved domains.

## Introduction

*Arabidopsis thaliana* genomic variation is available for investigation with the tools provided by the 1001 Genomes Project (1001genomes.org; 1001 Genomes Consortium, 2016). These tools help researchers filter their gene of interest for specific classes of variants (missense single nucleotide polymorphisms – SNPs, silent SNPs, insertions/deletions – indels, and truncations), and view different levels of variation across the 1001 Genomes data while overlaying genome annotation to give a holistic view of a variant’s possible impact. However, researchers are sometimes interested in comparing families of genes in *A. thaliana* by looking for differences in active sites or known functional domains. There is no simple online tool that shows variants across *A. thaliana* gene family alignments and plots population wide variation simultaneously: it is largely a process that must be done using desktop applications or command line scripts unfamiliar to biologists. Another level of complexity often not captured is that of alternative splicing: if investigators wished to explore variation in alternatively spliced genes they would need to compare the correct gene isoforms to fully understand what a given variant’s impact might be in a specific ecotype. A slightly different tool that allows a researcher to quickly overlay *A. thaliana* genes / gene products with additional annotation, such as that from the Protein Families Database (PFAM; Finn *et al*., 2014) and Conserved Domains Database (CDD; Marchler-Bauer *et al*., 2018) as presented in the Bio-Analytic Resource (BAR) ePlant Molecule Viewer (Waese *et al*., 2017), would assist researchers in obtaining a general overview of the variation in their genes of interest in the context of functional regions.

To address this resource gap, we developed Variant Viewer (http://bar.utoronto.ca/VariantViewer/) to investigate the 1001 Genomes data set, which allows researchers to view population level variation across splice isoforms, and across alignments of coding sequences of gene family members, with a useful level of annotation for orienting oneself. This web application is intended to act as a simple means for investigators to get a broad idea of the variation present in their gene / genes of interest, without having to navigate to several different data sources to collate the information on their own, and then to move on with further analyses based on easily-downloaded Variant Viewer outputs.

## Results

### The Variant Viewer

The Variant Viewer takes Arabidopsis Genome Initiative (AGI) identifiers as input and requests polymorphism information from the 1001 Genomes application programming interface (API). The response is used to prepare a visual summary of SNPs and indels along isoform gene model tracks (**Figure 1**). The coding sequences of known splice isoforms from the one or more inputted genes are then aligned and visualized with the frequencies non-synonymous SNPs. The graph also displays the location of CDD/PFAM domains and provides a colour coding for the physico-chemical properties of individual amino acids of each gene product. Alignment details are also available, specifying the actual aligned amino acids and highlighting the most prevalent variant at each site where applicable (**Figure 2**). The retrieved polymorphism data and alignment information can then be exported for further manipulation or analysis in JSON, CSV, or FASTA formats.

**Figure 1.**
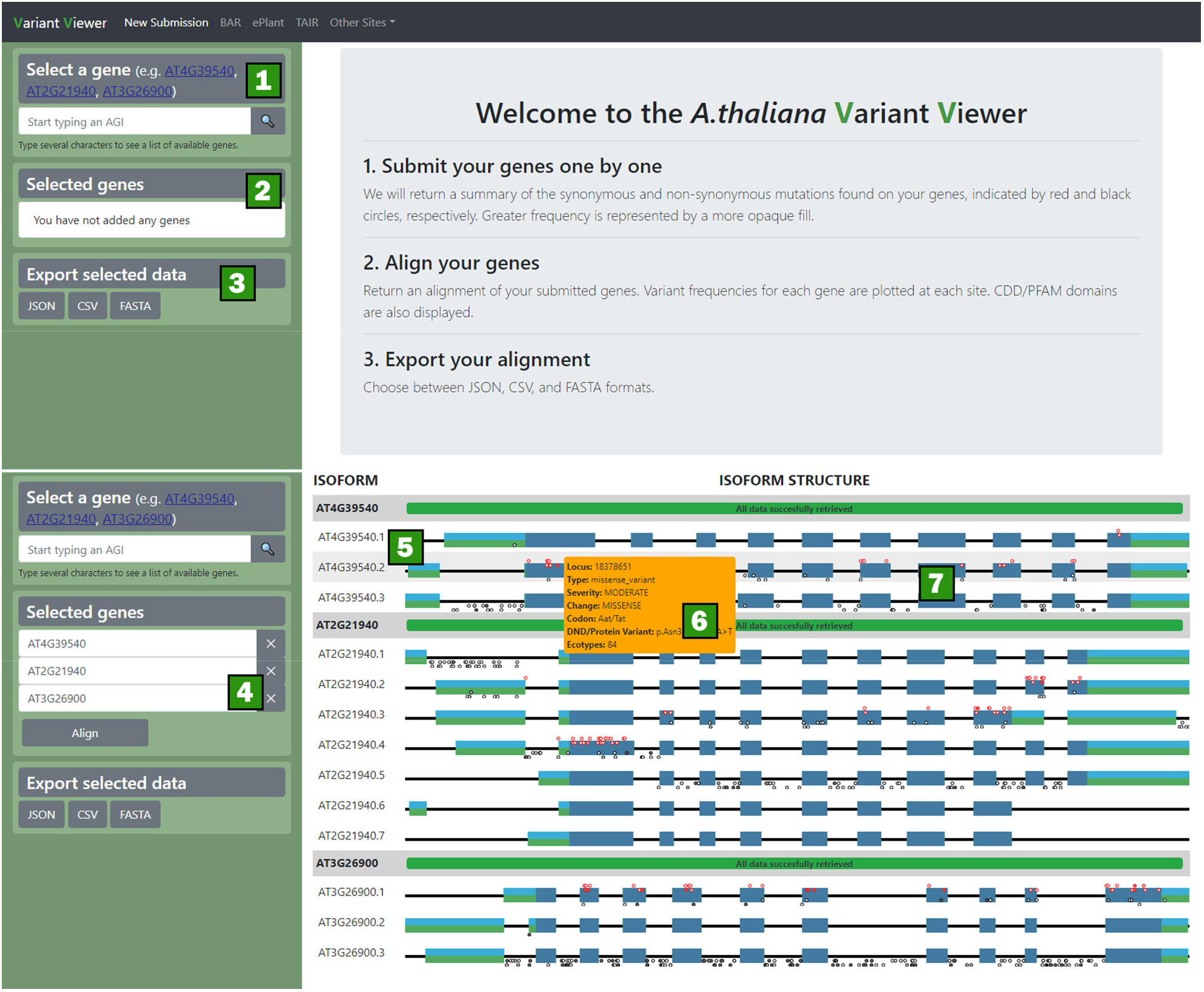
The Variant Viewer: an application for viewing variants from the 1001 Genomes Project. Graphics are rendered using d3.js and plotly.js (see Experimental Procedures). Top: landing page, bottom: output page. Variants are pulled on the fly from the 1001 Genomes Project API (http://tools.1001genomes.org/api) for each gene entered in the search box 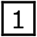. Selected genes 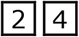 are displayed as simple gene models 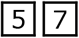 and variants are plotted onto models to give an overview of the variants in the gene region. Hover events are triggered allowing users to inspect individual exons, UTRs, coding sequences, and variants 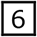. Users are also able to export data for their records 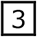.

**Figure 2.**
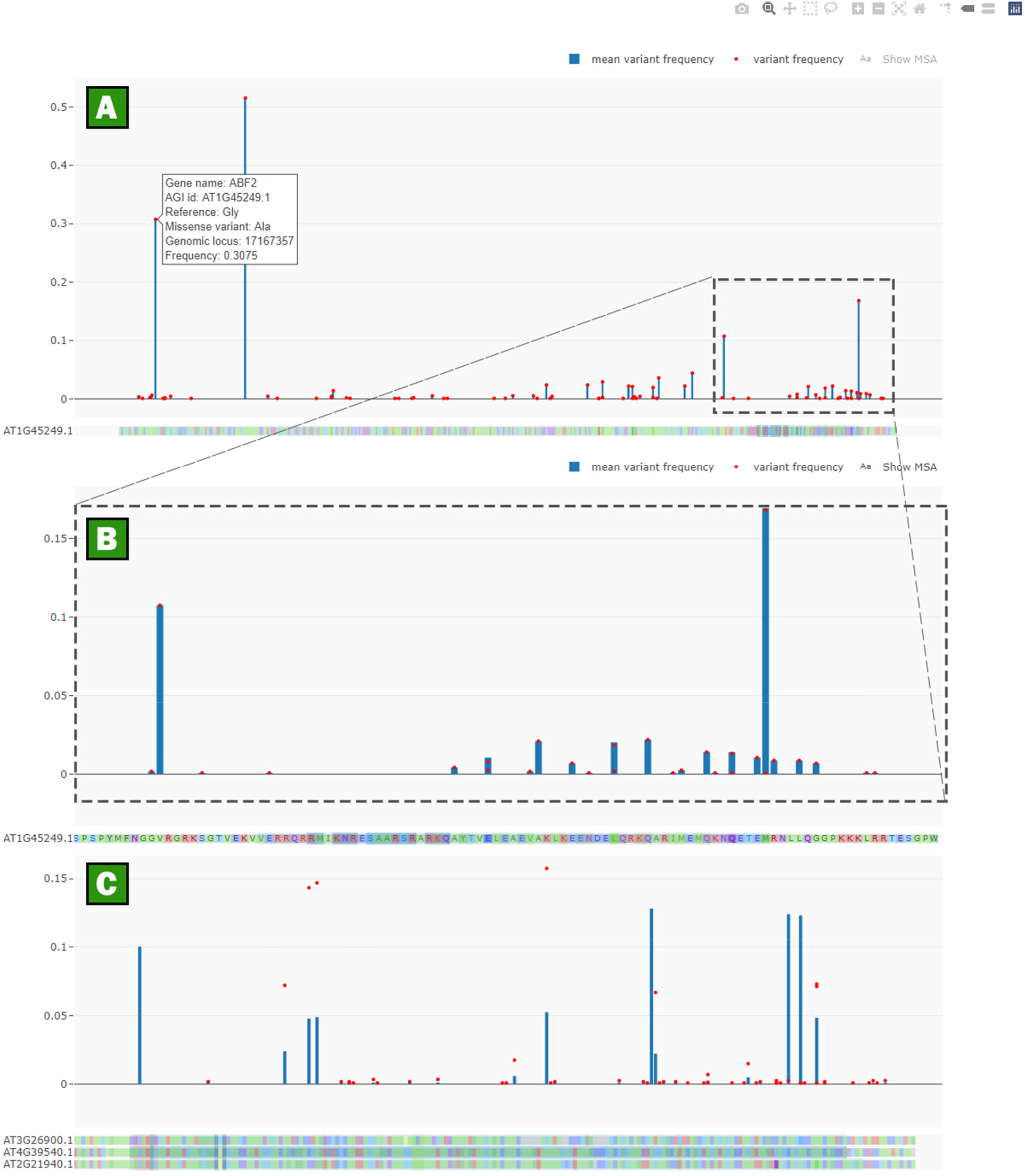
Graphs of protein sequences, MSAs of genes of interest, and variant frequencies. **A**. Individual variants are plotted as red dots for ABF2 (At1g45249), while mean variant frequency is shown by the blue bars, with the lower track showing the returned CDD domains for ABF2, namely the bZIP domain (medium blue background to track). Hover events are triggered for each point in the scatter plot allowing users to inspect individual variants. **B**. When researchers zoom in on the bZIP domain they can toggle to display the MSA and can see the known residues that govern dimerization between bZIP domains of ABF2 and the variants that are nearby. **C**. A multiple sequence alignment of SK1, SK2, and SKL1 shows highlighted residues corresponding to the shikimate binding, ADP binding sites, and magnesium binding domains as retrieved from the CDD.

### Natural variation in the ABA signaling members

As a use case for the Variant Viewer, we decided to compare the variation present in the “ core” ABA signaling pathway members. The variants present in the members of the ABA Response Element Binding Factors (ABFs), SnRK2s (Sucrose Non-Fermenting 1 Related Protein Kinase 2s), Group A Protein Phosphatase 2C (PP2Cs), and PYRABACTIN RESISTANCE1/PYR1-LIKE/REGULATORY COMPONENTS OF ABA RECEPTORS (PYR/PYL/RCARs) were retrieved from the 1001 Genomes Project site using Polymorph 1001 (https://tools.1001genomes.org/polymorph/). The PANTHER database (http://www.pantherdb.org; Mi *et al*., 2012) was queried for *A. thaliana* genes with annotations for abiotic stress-related genes, kinases, phosphatases, receptors, and transcription factors. AGI IDs were extracted from the retrieved lists and the missense variants were retrieved for each ID in the list. A boxplot of the variants organized by GO annotations reveals a lower number of SNPs in the core ABA signaling members than in the wider sets of proteins sharing similar annotations (**Figure 3**). However, a closer inspection of the individual proteins shows that PP2C phosphatases and the SnRK2 Ser/Thr kinases (blue points within the sets of phosphatases and Ser/Thr kinases, Figure 3) have slightly higher and lower numbers of variants, respectively, than genes in the broader annotations sets, suggesting that each signaling hub in the ABA core pathway might be evolving antagonistically compared to genes with similar GO annotations, and pointing to the utility of having a family-wide view of variation.

**Figure 3.**
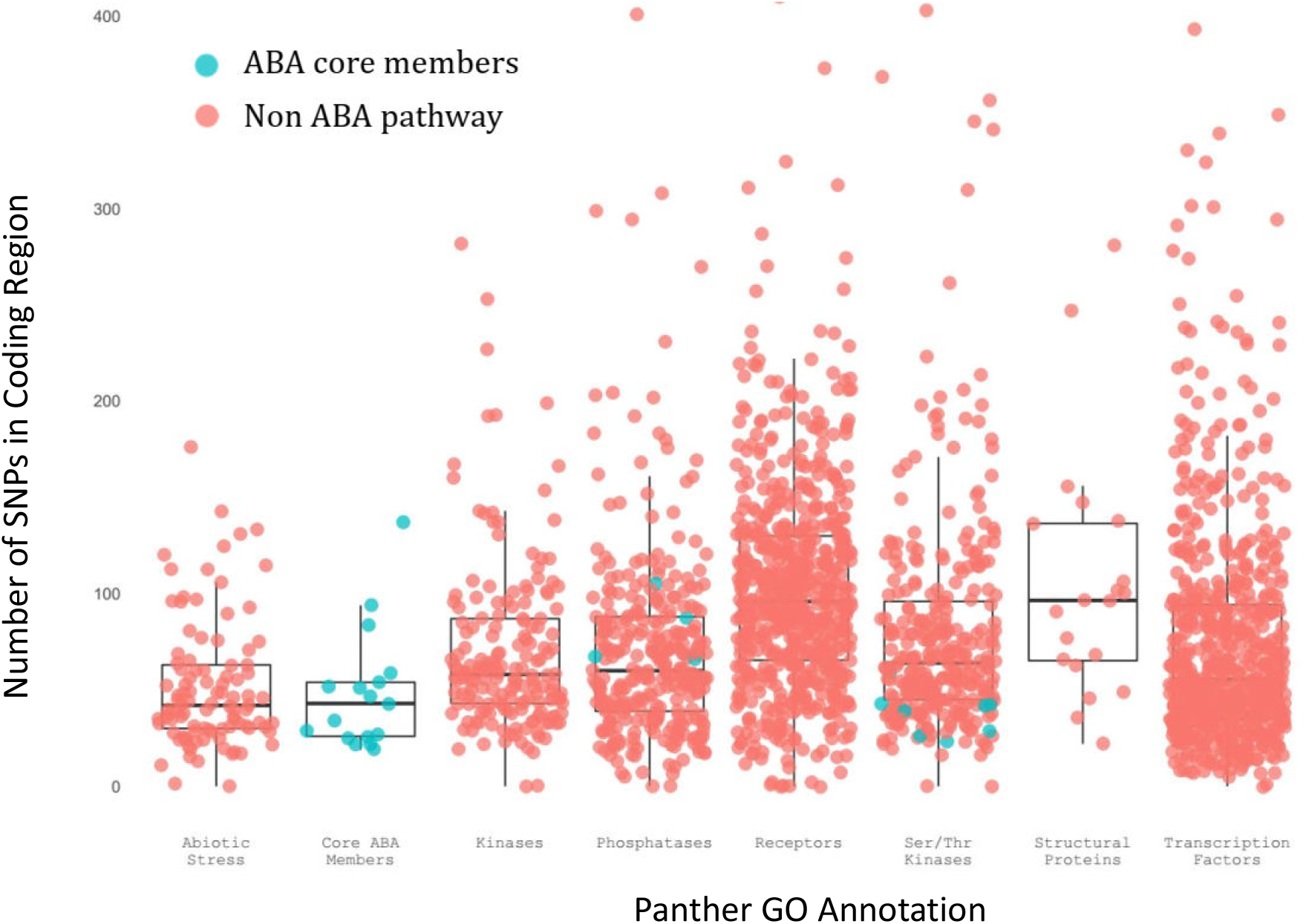
SNPs in the coding regions of *A. thaliana* genes plotted by PANTHER GO annotation. A general set of abiotic stress proteins and “ Core ABA Members” (highlighted in blue) are shown in addition to genome-wide sets of kinases, phosphatases, receptors etc. The abiotic stress and core ABA members have accumulated a median of approximately ∼45 missense variants per protein, slightly lower than the ∼52 missense variants per protein genome wide, but lower than the median for the broader sets of each protein functional class.

To compare potential variants contained within the ABF family the four representative isoforms of ABF1, ABF2, ABF3, ABF4 (AT1G49720.2, AT1G45249.3, AT4G34000.1, AT3G19290.3 respectively) were aligned using the Variant Viewer (**Figure 4**). Missense variants and non-synonymous variants largely occur across the entirety of the ABFs, but are totally absent within the DNA binding domain and at key leucine zipper residues in their basic leucine zipper (bZIP) domains (Jakoby *et al*., 2002). This suggests that these key residues are conserved across ecotypes and ABF family members and are thus the required residues for these bZIP transcription factors to function.

**Figure 4.**
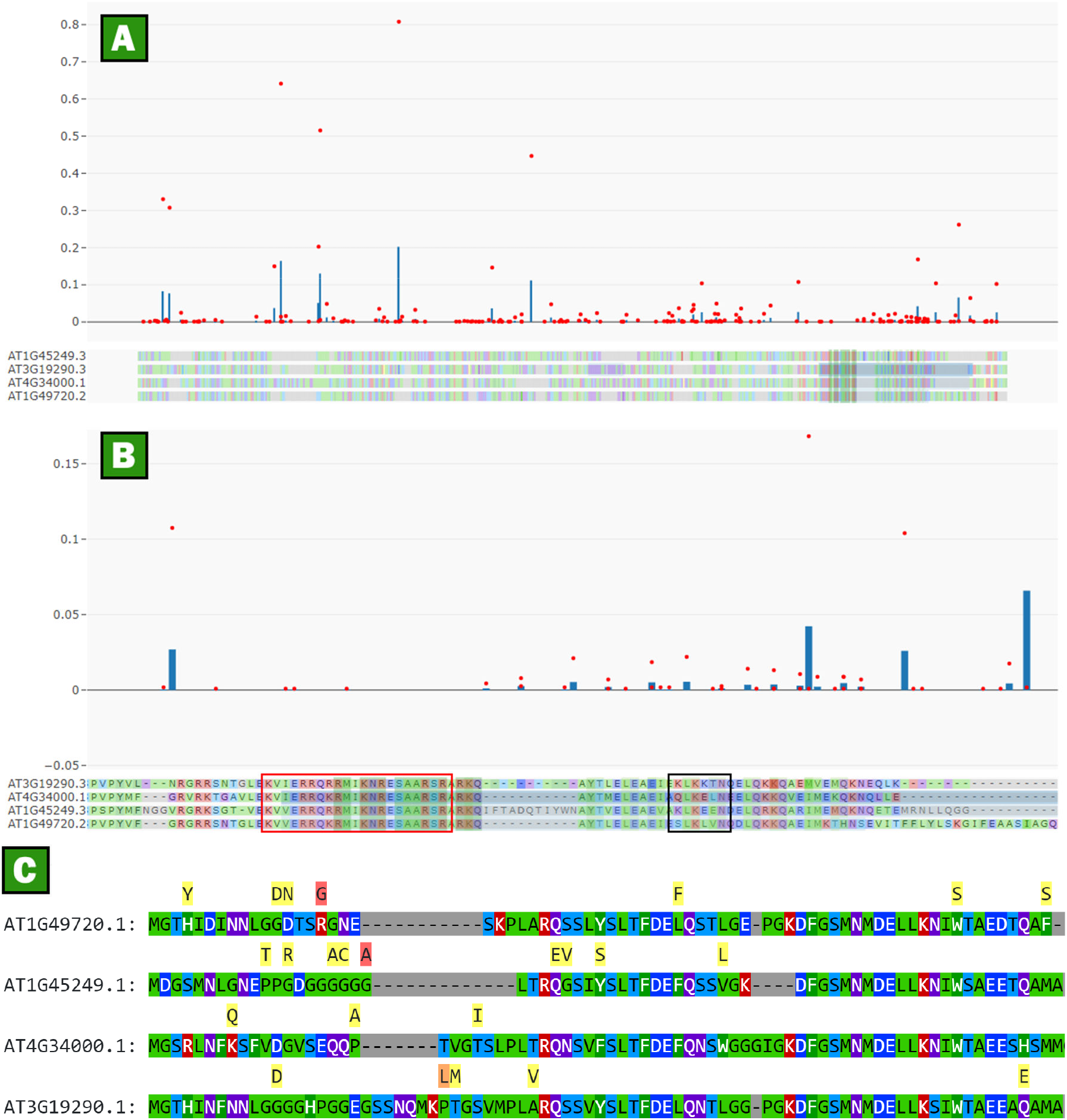
An alignment of the reference isoforms of the ABFs (ABF1/ABF2/ABF3/ABF4) with their associated variants. **A**. An overview of an alignment of the aligned ABFs (ABF1: AT1G49720.2; ABF2: AT1G45249.3; ABF3: AT4G34000.1; ABF4: AT3G19290.3). **B**. A close-up of C-terminal region of the ABFs shows few missense variants in the DNA binding domain (red box) compared to the rest of the protein sequence. The black box shows the dimerization region: missense variants are present in between critical dimerization residues of the bZIP domain but do not occur at any critical hydrophobic residues. **C**. Clicking on the “ Alignment Details” output in the Variant Viewer app calls up an alignment – shown here with different isoforms of the ABFs – with additional tracks denoting the most frequent non-synonymous change at a given location superposed above the protein sequences, with background colouring denoting the frequency of the substitution (red = more frequent, yellow = less frequent).

Two residues are contained within the bZIP domain but do not fall directly on the essential leucine dimerization residues or others noted to be involved in dimerization (Vinson *et al*., 1993). There are no missense variants that directly overlap the conserved RxxS/T phosphorylation sites in the N terminus of the protein, but there is a low frequency (in just 5 ecotypes) early stop codon in ABF2 at site 62 (Q to *), which would result in a 61-residue truncated ABF2 protein. Coincidentally, truncations of ABF2 at site 62 have been previously tested in cultured T87 *A. thaliana* protoplasts and have been shown to be able to strongly activate reporters when fused to a non-native DNA binding domain such as GAL4; however, the truncation should totally abolish the activity of ABF2 as it occurs early in the first exon of ABF2 and removes the bZIP domain in the C terminus of the protein. A second truncation in ABF2 is found at site 263 (Gly to *, in 3 ecotypes), which still places it before the bZIP domain, rendering ABF2 non-functional. Null mutants of ABF2 have no discernable growth or drought stress phenotype due to the overlapping redundancy of ABF2, ABF3 and ABF4 (Yoshida *et al*., 2010).

Additionally, the accessions containing these early truncations are all Swedish in origin and are found at a higher latitude where drought may be less common. However, ABF2 has more specific localization to roots, has an essential role in glucose signaling and in regulating early seedling development by driving the expression of hexokinases which initiate glycolysis (Kim *et al*., 2004), and has a narrower role in osmotic stress responses alongside ABF3. It would be interesting to investigate variants in proteins that complement the truncated ABF2s in those ecotypes to see if those populations are modulating these responses in a novel way to account for the loss of ABF2, for example, ABF3 in terms of osmotic stress, or other transcription factors that drive the expression of glucose metabolism genes. In this way an investigator might uncover a locally adapted response to a novel variant that would otherwise go unnoticed if one only considered the genes in the Col-0 ecotype.

## Discussion and future directions

Although many of the basic features of the Variant Viewer app have been implemented, it still has room for improvement. Implementing a pane suggesting homologs of the query gene would make investigating variation in gene family members easier. Giving users the option to upload their own MSAs would provide them more control over how to generate their MSAs, compared to how the Variant Viewer currently uses default values for DECIPHER in R (Wright, 2016) to align sequences. The “ export output” could also be more intuitively formatted for downstream analyses.

Additionally, although Variant Viewer has undergone some user testing (and in fact was critiqued by 10 students as part of a user experience module in a graduate-level course on data visualization), we expect to fix minor bugs and address edge cases on an *ad hoc* basis in response to further user feedback. Nevertheless, this tool helps researchers to “ think big” regarding their possible experiments, and to take advantage of the useful 1001 Genomes resources for possible hypothesis generation.

## Experimental procedures

### Data sources and Javascript libraries

To construct gene models, and plot variants, the tool requests data from several sources when a researcher enters an Arabidopsis Genome Initiative identifier (AGI ID). First the Araport11 gene structure for the AGI ID is requested from a BAR web service (http://bar.utoronto.ca/webservices/bar_araport/gene_summary_by_locus.php?locus=AGI_ID), which returns genomic positions for each exon, intron, and untranslated region for gene models associated with the requested AGI ID. Next, another BAR web service (http://bar.utoronto.ca/webservices/bar_araport/get_protein_sequence_by_identifier.php?locus=AGI_ID) returns the protein sequence on an isoform by isoform basis, followed by a request to the 1001 Genomes application programming interface (API) (http://tools.1001genomes.org/api/v1.1/effects.json?type=snps;accs=all;gid=AGI_ID) requesting Variant Call Format data annotated by SNPeff (Cingolani *et al*., 2012) for the selected AGI ID. All AGI ID isoforms that have valid gene models have corresponding protein sequences (for example AT1G45249 returns 10 gene isoforms, and AT1G45249.12 does not a corresponding protein sequence), but not all gene isoforms have corresponding variant entries in the 1001 Genomes database. This is because the variants are assigned to the transcript that they map most closely to instead of to the representative gene model (for example AT1G45249 isoforms .1, .2, and .3 all have mapped variants but only AT1G45249.3 is the representative model). If the 1001 Genomes variant query is successful, a final set of requests are made to two BAR webservices (http://bar.utoronto.ca/eplant/cgi-bin/CDDannot.cgi, http://bar.utoronto.ca/eplant/cgi-bin/PfamAnnot.cgi) that return PFAM (Finn *et al*., 2014) and CDD (Marchler-Bauer *et al*., 2018) domains for the AGI ID’s protein sequence. CDD requests return a domain annotation and individual residues for a location in a protein query (for example “ DNA binding domain: K235, L238, I241”), whereas PFAM requests return a domain, annotation, expectation score and start / end locations (for example “ DNA Binding Domain: [Start: 235, End: 241, Expect 2E-45, Annot: Basic Leucine Zipper]”). It should be noted that since different isoforms of genes have different coding sequences, and each is submitted separately as a query, some isoforms may return domains while others may not. All returned data are in JSON format and are saved to objects that can be exported by researchers after submitting their query if they wish to have the original information for reference or further use.

The application interface is built using “ vanilla” Javascript, and a Bootstrap 4 (https://getbootstrap.com) template and style layout. Graphics and tables are added to the web app’s “ document object model” using d3.js (https://d3js.org; Bostock *et al*., 2011). The variant and coding sequence plot is rendered using the open source plotly.js (https://plot.ly/javascript; Sievert, 2020) library which allows the user to interact with the convenient plotly.js interface (zooming, export, point selection etc.).

The web application is split into five main components, a controller script that determines what is being displayed to the user (main.js), a data script that asynchronously requests and formats data (varData.js), two plotting scripts that render images into graphics for the user (plots.js and overview.js), and a backend R script (R-Development-Core-Team, 2008) that aligns proteins and maps their locations (alignProteins.R). The scripts can be accessed by looking at github account associated with the project (https://github.com/daea/Variant_viewer).

### Application logic

Upon navigating to the Variant Viewer the user is greeted with a simple Bootstrap 4 responsive layout and a splash page that provides instructions on how to use the app and sample genes (Figure 1). The title bar provides links to several databases and viewers for information on their gene of interest. The side input panel (Figure 1, Item 1) contains a dropdown suggestion search box that will autocomplete AGI ID and gene names and restrict the user to entering only valid gene IDs. When users have entered a valid AGI ID they click the “ Add Gene” button to select a gene and request relevant data from the data sources mentioned above. If an improper AGI ID is entered the application will prompt users with an alert box and they will be given a chance to re-enter an AGI ID. The “ Selected genes” panel (Figure 1, Items 2 and 4) is then populated with that gene and the gene models are drawn in the main app panel to the right.

Underneath the Selected genes panel is an export panel that allows users to export their data if desired (Figure 1, Item 3). As users “ Add genes” to their selected genes list, the main panel draws the respective gene isoforms and sends a request to the 1001 Genomes API for the variant data (Figure 1, Item 5). The gene models are scalable vector graphics (SVG) images that dynamically scale to fill the page width when they are rendered. Gene models are plotted in three colours to represent exons, untranslated regions and coding regions (Figure 1, Item 7). All isoforms of a gene model are scaled to the total length of a gene on the chromosome, so the different features line up and allow the user to easily see which features are retained in alternative splice variants. The user can hover over the gene models to view chromosomal position for the type of feature they are interested in (Figure 1, Item 6).

When the 1001 Genomes data have been retrieved and parsed, the application plots variants onto the gene models. The positions of variants on the horizontal are calculated relative to the width of the parent gene model to reflect the genomic position of a variant in the gene. Variants are currently plotted in two colours, red and black, for missense variants and all other variants respectively. The shading of the circle fill (red or black) currently reflects the number of ecotypes (out of 1135) that contain the respective variant. Similar to the gene models, when users hover over the variants they are provided with a tooltip that displays information about that variant such as the positions, severity, type, codon change (if applicable), the variant code as called by SNPeff and the number of ecotypes that contain the variant. If a user selects multiple genes, new requests are made, and gene models are rendered for each gene.

After users have selected several genes, if desired, they may submit the genes for alignment. At this point the AGI IDs and protein sequences are submitted to a server-side PHP script that passes the data to an R script which performs a multiple sequence alignment (MSA) using the DECIPHER function AlignSeqs (Wright, 2016). The R script then returns a matrix that relates the new position in the MSA to the actual positions of the amino acids in their coding sequences. This matrix is then used to map the variant positions in the requested 1001 Genomes data to the multiple sequence alignment (for example in an alignment of two proteins position 3 in an original CDS may correspond to position 8 in the MSA due to the insertion of gaps prior to position 3). The application then renders a plotly.js scatter plot above the table of gene models which shows the number of ecotypes containing a given variant relative to their position in the multiple sequence alignment (Figure 2). The bottom subplot displays amino acid colour-coding, and PFAM and CDD domains corresponding to each of the plotted genes as coloured boxes overlaying the MSA. Additionally, the MSA can be displayed by clicking the “ Show MSA” entry in the figure legend, which then displays the residues for each aligned coding sequence (Figure 2C). The user may zoom in on the plot and highlight variants by selecting them, export the graph as a portable network graphics (PNG) file or edit the graph using the online plotly.js interface (Figure 2B). Additionally, by clicking on legend entries a user can choose to remove missense, synonymous or premature stop codons to filter the displayed data as desired.

A text-based visualization of the alignment is also provided, hidden in a modal that is revealed when “ Alignment Details” is selected. The specific amino acid variant is displayed above its alignment site, and is coloured based on its frequency. Red indicates frequencies equal to or greater than 10% (found in at least 113/1135 ecotypes), yellow for those equal to or less than 0.1% (found in only 1/1135 ecotypes), and orange for those at an intermediate frequency to those values.

## Acknowledgements, Funding and Author Contributions

We are grateful to Dr. Barb Moffatt from the University of Waterloo for providing feedback on an initial version of Variant Viewer. NJP was funded by an NSERC Discovery Grant. MC and EE developed the Variant Viewer code and drafted the manuscript. VL organized user testing and provided manuscript and interface feedback. AP implemented webservices and APIs at the BAR to power the Variant Viewer front end. NJP supervised all co-authors and edited the final version of the manuscript.

## References

1001 Genomes Consortium (2016) 1,135 Genomes Reveal the Global Pattern of Polymorphism in Arabidopsis thaliana. Cell,166, 481–491.

Bostock, M., Ogievetsky, V. and Heer, J. (2011) D3 Data-Driven Documents. IEEE Trans. Vis. Comput. Graph.,17, 2301–2309.

Cingolani, P., Platts, A., Wang, L.L., Coon, M., Nguyen, T., Wang, L., Land, S.J., Lu, X. and Ruden, D.M. (2012) A program for annotating and predicting the effects of single nucleotide polymorphisms, SnpEff. Fly (Austin),6, 80–92.

Finn, R.D., Bateman, A., Clements, J., et al. (2014) Pfam: The protein families database. Nucleic Acids Res.,42, 222–230.

Jakoby, M., Weisshaar, B., Dröge-Laser, W., Vicente-Carbajosa, J., Tiedemann, J., Kroj, T. and Parcy, F. (2002) bZIP transcription factors in Arabidopsis. Trends Plant Sci.,7, 106–111.

Kim, S., Kang, J.Y., Cho, D.I., Park, J.H. and Soo, Y.K. (2004) ABF2, an ABRE-binding bZIP factor, is an essential component of glucose signaling and its overexpression affects multiple stress tolerance. Plant J.,40, 75–87.

Marchler-Bauer, A., Derbyshire, M.K., Gonzales, N.R., et al. (2018) CDD : NCBI ‘ s conserved domain database., 43, 222–226.

Mi, H., Muruganujan, A. and Thomas, P.D. (2012) PANTHER in 2013: modeling the evolution of gene function, and other gene attributes, in the context of phylogenetic trees. Nucleic Acids Res.,41, D377–D386.

R-Development-Core-Team (2008) R: A language and environment for statistical computing.

Sievert, C. (2020) Interactive Web-Based Data Visualization with R, plotly, and shiny, Chapman and Hall/CRC. Available at: https://plotly-r.com.

Vinson, C.R., Hai, T. and Boyd, S.M. (1993) Dimerization specificity of the leucine zipper-containing bZIP motif on DNA binding: Prediction and rational design. Genes Dev.,7, 1047–1058.

Waese, J., Fan, J., Pasha, A., et al. (2017) ePlant: Visualizing and Exploring Multiple Levels of Data for Hypothesis Generation in Plant Biology. Plant Cell,29, 1806– 1821.

Wright, E.S. (2016) Using DECIPHER v2. 0 to Analyze Big Biological Sequence Data in R., 8, 352–359.

Yoshida, T., Fujita, Y., Sayama, H., Kidokoro, S., Maruyama, K., Mizoi, J., Shinozaki, K. and Yamaguchi-Shinozaki, K. (2010) AREB1, AREB2, and ABF3 are master transcription factors that cooperatively regulate ABRE-dependent ABA signaling involved in drought stress tolerance and require ABA for full activation. Plant J.,61, 672–685.

